# Sensory evidence accumulation using optic flow in a naturalistic navigation task

**DOI:** 10.1101/2021.04.26.441532

**Authors:** Panos Alefantis, Kaushik J. Lakshminarasimhan, Eric Avila, Jean-Paul Noel, Xaq Pitkow, Dora E. Angelaki

## Abstract

Sensory evidence accumulation is considered a hallmark of decision-making in noisy environments. Integration of sensory inputs has been traditionally studied using passive stimuli, segregating perception from action. Lessons learned from this approach, however, may not generalize to ethological behaviors like navigation, where there is an active interplay between perception and action. We designed a sensory-based sequential decision task in virtual reality in which humans and monkeys navigated to a memorized location by integrating optic flow generated by their own joystick movements. A major challenge in such closed-loop tasks is that subjects’ actions will determine future sensory input, causing ambiguity about whether they rely on sensory input rather than expectations based solely on a learned model of the dynamics. To test whether subjects performed sensory integration, we used three independent experimental manipulations: unpredictable optic flow perturbations, which pushed subjects off their trajectory; gain manipulation of the joystick controller, which changed the consequences of actions; and manipulation of the optic flow density, which changed the reliability of sensory evidence. Our results suggest that both macaques and humans relied heavily on optic flow, thereby demonstrating a critical role for sensory evidence accumulation during naturalistic action-perception closed-loop tasks.

## Introduction

To survive in a perpetually uncertain and volatile world, we must make sequential decisions within a limited time horizon. To succeed, we accumulate information from our noisy environment to inform decisions that may lead to desirable outcomes. Sensory evidence accumulation is considered a hallmark of perceptual decisions and is used to reduce uncertainty in favor of the optimal potential action. However, insights about how the brain integrates sensory evidence come largely from simple laboratory tasks in which sensory cues are discrete (e.g. auditory clicks) and/or have statistics that remain stationary over time (e.g. random dot kinematograms) (de Bruyn & Orban, 1988; Glass & Pérez, 1973; Snowden & Braddick, 1990; Watanabe & Kikuchi, 2006; de Lafuente et al., 2015; Gold & Shadlen, 2000; Kim & Shadlen, 1999; Liu & Pleskac, 2011; Drugowitsch et al., 2015; Gu et al., 2008; Hou et al., 2018). In contrast, in the real world, the statistics of sensory inputs can change continuously depending on the actions we take, creating a closed-loop interaction between perception and action. To study computations that underlie dynamic, closed-loop behaviors, we need to employ tasks that closely resemble naturalistic environments with moderate complexity that strike a balance between recapitulating the rich dynamics of the world and exerting control over the task variables.

One example of real-world sequential action-perception interaction experiences is *path integration*, an ethological behavior that involves integrating optic flow cues, which are generated by one’s own self-motion (humans: Butler et al., 2010; Ellmore and McNaughton, 2004; Frenz and Lappe, 2005; Frenz et al., 2007; Kearns et al., 2002; Lappe et al., 2007; Wiener et al., 2016, insects: Collett and Collett, 2017, 2000; Heinze et al., 2018; Stone, 2017, rodents: (Campbell et al., 2018; Kautzky and Thurley, 2016; Thurley and Ayaz, 2017). Integrating self-motion cues helps maintain one’s sense of position, even when explicit position cues are unavailable (Collett and Collett, 2000; Etienne and Jeffery, 2004; Golledge, 1999). During rotational self-motion, for example, optic flow indicates a change in angular position. During translation, the radial pattern of optic flow provides information needed to estimate changes in displacement. Although optic flow-based path integration is a real-world behavior likely involving time integration of self-motion velocity cues, it is seldom exploited as a sensory evidence accumulation task.

Instead, primate studies of optic flow evidence accumulation have been limited to passive viewing laboratory tasks, where sensory cues and actions are discrete (e.g. in 2 alternative-forced-choice) and intermittent (e.g. end of trial) (Drugowitsch et al., 2015; Gu et al., 2008; Hou et al., 2018). This is also true for visual motion in general, a classical stimulus for sensory-based decision-making studies (Gold and Shadlen, 2007). Such laboratory tasks of sensory evidence accumulation differ strikingly from real life experiences rich in sequential actions with temporal dynamics, as well as continuous monitoring of the ceaseless stream of evidence impinging on sensory receptors. Do the principles learned from artificial laboratory tasks extend into naturalistic behaviors?

To be able to address this question, we recently developed a naturalistic visuomotor virtual navigation task where subjects use a joystick controller to navigate to a cued target location on the ground plane using optic flow cues. Previous work using this paradigm tested whether self-motion estimates are integrated optimally to compute position (Lakshminarasimhan et al., 2018a; Noel et al., 2020), and whether the resulting beliefs about position may be reflected in eye movements (Lakshminarasimhan et al., 2020). However, we do not know the extent to which self-motion estimates in this task originate from integrating optic flow. To exploit this task in the context of sensory-based sequential decision-making, we must first test whether subjects do integrate optic flow and rule out other navigation strategies that rely solely on an internal model of the control dynamics. Here we adjudicate between the above alternatives by employing three variations of this task. First, we incorporated optic flow perturbations in random directions and amplitudes to test subjects’ dynamic compensation. Second, we manipulated the gain of the joystick controller, challenging the subjects to adjust their actions to navigate to the desired location. Third, we manipulated the density of optic flow, thus varying the reliability of the sensory cues. We show that both macaques and humans rely on optic flow to perform this task, allowing for the study of neural correlates of sensory-based decision-making in naturalistic tasks akin to ethological dynamic behaviors.

## Results

Macaque and human subjects performed a visual navigation (‘firefly’) task in which they used a joystick to steer to a briefly cued target location in a virtual environment devoid of landmarks (**Fig. 1A**, see **Methods**). In each trial, a circular target appeared briefly on the ground plane at a random location within the subject’s field of view (**Fig. 1B**). Subjects had to navigate to the remembered target location using a joystick to control their linear and angular velocity. The task goal was to stop within the circular reward zone of the target. Unless stated otherwise, feedback was provided immediately after the end of each trial (**Fig. 1C**; Methods). The virtual ground plane elements were transient and could therefore not be used as landmarks—only to provide optic-flow information.

**Figure 1:**
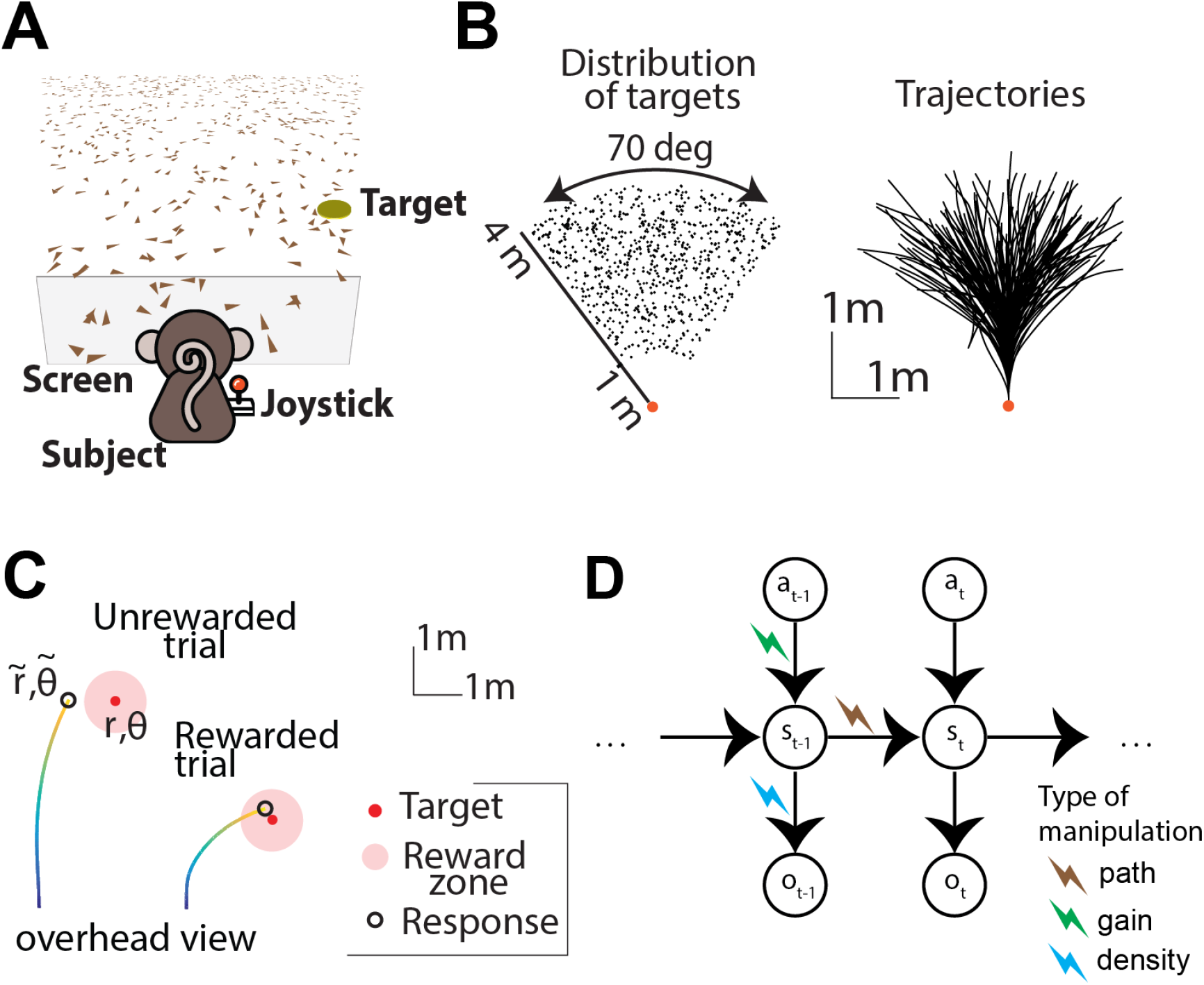
**(A)** Monkeys and humans use a joystick to navigate to a cued target (*yellow disc*) using optic flow cues generated by ground plane elements (*brown triangles*). The ground plane elements appeared transiently at random orientations to ensure that they cannot serve as spatial or angular landmarks. **(B)** *Left:* Overhead view of the spatial distribution of target positions across trials for monkey subjects. Positions were uniformly distributed within subjects’ field of view. The actual range of target distances and angles was larger for human subjects (**see methods**). *Right:* Movement trajectories of one monkey during a representative subset of trials. Orange dot denotes starting position. **(C)** Example trials showing incorrect (*left*) and correct (*right*) responses. Note that subjects had to stop within a ±0.6m zone to receive reward. **(D)** Schematic representation of the Markov decision process that governs self-motion sensation. a, s and o denote action (joystick input), state (velocity, position), and sensory observations, respectively, while subscripts denote time indices. We used 3 manipulations to alter the causal dependency of the variables involved in the navigation process: visual perturbations (brown bolt), gain manipulations (green bolt), density manipulation (blue bolt).

This active control paradigm can be understood as a Partially Observable Markov Decision Process (POMDP)(Åström, 1965; Kwon et al., 2020; Sutton and Barto, 1998) in which the sensory observation *o_t_* (optic flow) is determined by the current state *s_t_* (velocity), which depends only on the previous state *s*_*t*-1_ and the current action *a_t_* (joystick movement) through the control dynamics (**Fig. 1D**; Methods - **Eq 1**). To perform this task optimally, subjects must combine their knowledge of the control dynamics with the sensory observation to estimate their current velocity and integrate that estimate over time such that they can stop upon reaching the reward zone. In principle, however, subjects could choose to ignore sensory inputs and still perform reasonably well by dead reckoning with an accurate internal model of the control dynamics.

In separate sessions, we used three different manipulations of this ‘firefly’ task to test whether subjects used sensory evidence accumulation, i.e., integrated optic flow. The causal effect of these manipulations on the decision process is illustrated in **Fig. 1D**: (i) Random perturbations, which imposed an external passive displacement that moved the subjects away from their expected path, disrupting the transition to the desired state (**Fig. 1D**, red bolt); (ii) Altered gain of the joystick controller from that used during training changed the effect actions induced upon the current state (**Fig. 1D**, green bolt); (iii) Different densities of the ground plane elements manipulated the reliability of the observations provided by each state (**Fig. 1D**, blue bolt). Next, we explore steering responses for each of these experimental manipulations.

### Subjects compensate for unpredictable perturbations

In a random half of the trials, subjects were gradually displaced from their controlled trajectory (visual trajectory perturbation) while steering, challenging them to counter the displacement to reach their goal location. The perturbation began after a random delay (0-1 s) following the subject’s movement onset and consisted of independent linear and angular components whose velocity profiles followed a Gaussian (monkeys) or triangular (humans) waveform (see Methods) lasting one second. The onset delay and the amplitude of each perturbation were drawn randomly from uniform distributions (**Fig. 2A**). To reach the target, subjects needed to update their position estimates based on the amplitude and direction of the imposed perturbation, which could only be estimated by sensory integration of visual motion cues (optic flow).

**Figure 2:**
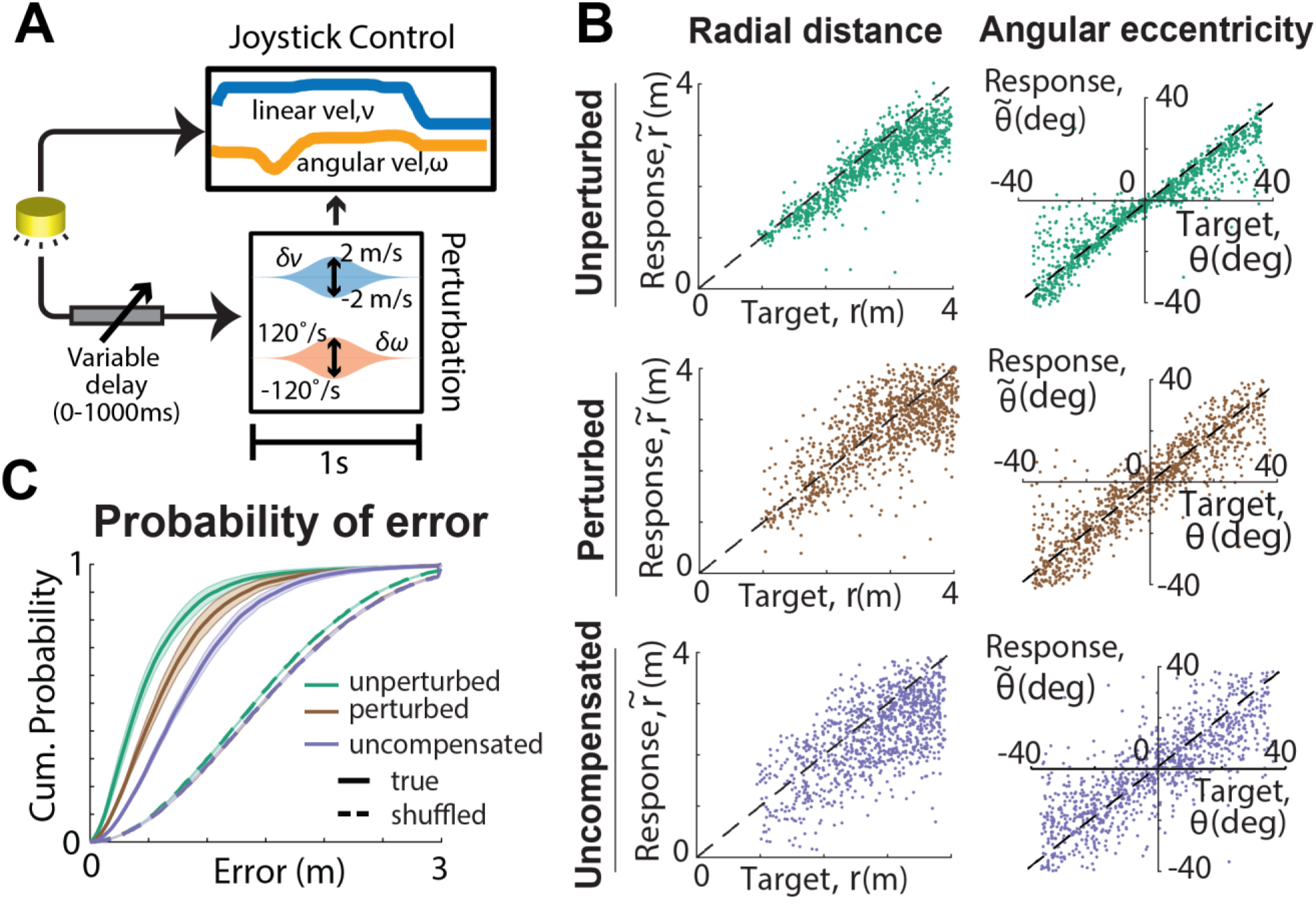
**(A)** Schematic illustration of the perturbation manipulation: after a variable delay (0-1s) from movement onset, a fixed duration (1s) perturbation was added to the subjects’ instantaneous self-motion velocity. Perturbations consisted of linear and angular velocity components with Gaussian profiles whose amplitudes varied randomly across trialsJ from −2 to 2m/s and from −120 to 120°/s for the linear and angular velocities, respectively. **(B) Left:** Comparison of the radial distance 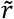 of an example subject’s response (final position) against the radial distance *r* of the target for unperturbed (top, green), perturbed (middle, brown), and the simulated uncompensated case (bottom, purple) trials. **Right**: Similar comparison for the angular eccentricity of the same subject’s response 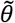 versus target angle *θ*. Black dashed lines show unity slope. **(C)** Cumulative probability of error magnitude for unperturbed (*green*), perturbed (*brown*) and simulated (uncompensated case; *purple*) trials, averaged across all monkeys. *Solid lines* and *dashed lines* represent the values obtained from the actual and shuffled data, respectively. Shaded regions represent ±1 SEM.

We compared the subjects’ responses (i.e., stopping location) in each trial to the corresponding target location separately for unperturbed and perturbed trials. We also simulated responses for a ‘uncompensated’ case where subjects steer towards the original target, completely ignoring imposed perturbations. We generated the uncompensated responses by adding the linear and angular velocities of each perturbation to the self-motion velocity profiles of the monkeys in target-matched trials without perturbations (see **Methods, Equation 2**). For each condition (unperturbed, perturbed, uncompensated), we calculated the radial distance 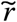 and angular eccentricity 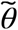 of the subjects’ final position, which were then compared to the initial target distance *r* and angle *θ*. Monkeys behaved well on this task, steering appropriately toward the targets in perturbed and unperturbed conditions. As shown with an example monkey session in **Fig. 2B** both radial distance and angular eccentricity of the monkeys’ responses were highly sensitive to target location for both unperturbed (Pearson’s *r* ± SD, radial: 0.73 ± 0.1, angular: 0.91 ± 0.03) and perturbed (radial: 0.59 ± 0.08, angular: 0.85 ± 0.05) trials. Furthermore, although perturbations decreased the correlation compared to unperturbed trials for both radial distance (*t*-test, *p* < 10^-6^) and angular eccentricity (*t*-test, *p* < 10^-6^), this decline was less than what would be expected from the uncompensated case (radial: 0.56 ± 0.06, *p* = 0.004, angular: 0.76 ± 0.05, *p* < 10^-6^). Results in humans were qualitatively similar (**Suppl. Table1**).

To more directly test whether subjects compensated for the perturbations, we computed the absolute error – the distance between the stopping position and the target – on each trial. In monkeys, errors in trials with perturbations (0.71 ± 0.08 m s.d.) were larger (*p* < 10^-6^) than those without perturbations (0.55 ± 0.07m), but smaller than the uncompensated case (0.87 ± 0.08 m, *p* < 10^-6^). In humans, steering accuracy in perturbation trials were not significantly different than in non-perturbation trials, and significantly better than the uncompensated condition (3.19 ± 1.61m (SD) vs. 3.13 ± 1.86m w/o perturbations, *p* = 0.61; uncompensated case: 4.21 ± 1.78 m, *p* < 10^-6^). Thus, perturbations decreased steering accuracy relative to unperturbed trials in monkeys (but not humans), but this increase was much less than expected from the uncompensated case (**Fig. 2C**).

To quantify performance accuracy across humans and monkeys, we adopted the approach of receiver operating characteristic (ROC) to continuous responses. Specifically, for each subject, we computed the actual reward rate and the chance level reward rate, obtained by shuffling target locations across trials, as a function of a hypothetical reward window size (Lakshminarasimhan et al., 2020). We obtained the ROC curves by plotting the actual responses against the responses at chance level and computed the area under the ROC curve (AUC) (**Fig. 3A**). Chance performance would be reflected by an AUC of 0.5, while perfectly accurate performance would yield an AUC of 1. We compared the area under the curve across conditions for monkeys (mean ± SD, unperturbed: 0.87 ± 0.03, perturbed: 0.83 ± 0.08, uncompensated case: 0.76 ± 0.03) and humans (unperturbed: 0.75 ± 0.03, perturbed: 0.72 ± 0.08, uncompensated case: 0.61 ± 0.08). Although the perturbations reduced response accuracy relative to the unperturbed trials (t-test: *p*= 0.003), this reduction was much less than expected for uncompensated perturbations (*p* < 10^-6^) (**Fig. 3B**). These results show that subjects were able to compensate for optic flow perturbations, supporting the hypothesis that they integrate optic flow for path integration.

**Figure 3:**
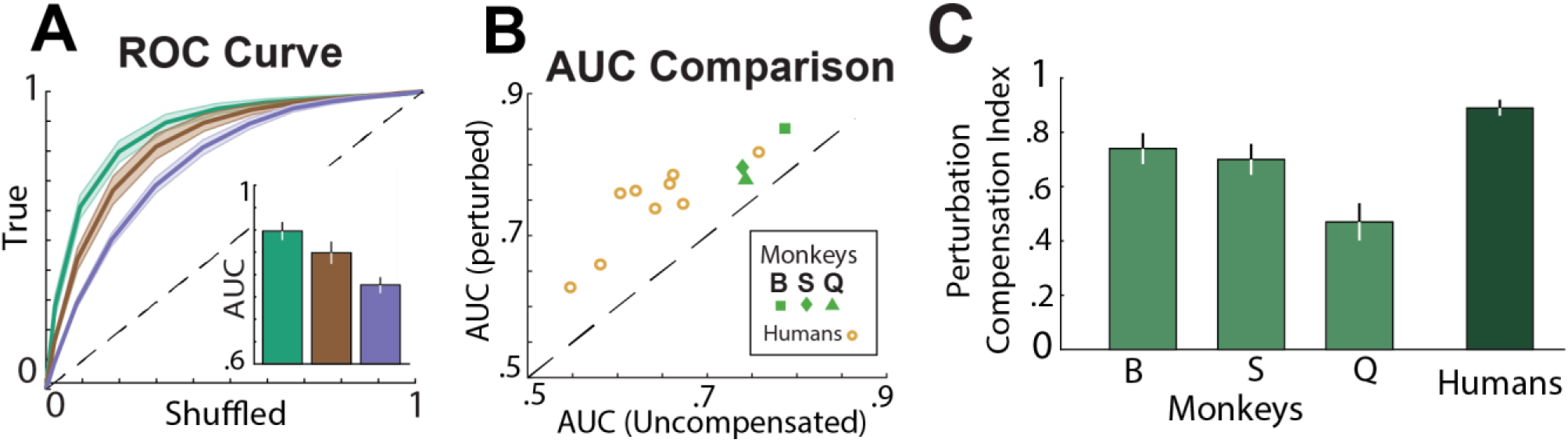
**(A)** ROC curves for unperturbed (*green*), perturbed (*brown*) and simulated (uncompensated case; *purple*) trials, averaged across all monkeys. **Inset in A:** Area under the curve (AUC) for the corresponding conditions in **(A)**. Shaded regions and error bars denote ±1 SEM. **(B)** Comparison of the area under the curve (AUC) of the perturbation (*true*)and simulated uncompensated trials separately for each monkey and human. **(C)** Bar plot of the mean perturbation compensation index for individual monkeys (*light green*) and average human subjects (*dark green*). Error bars denote ±1 SEM.

To further quantify the extent to which the subjects compensated for perturbations, we computed a perturbation compensation index (PCI; see **Methods**). A value of 0 denotes an accuracy equal to the uncompensated case response and thus a complete lack of compensation, whereas a value of 1 denotes an accuracy equivalent to the unperturbed trials and thus represents perfect compensation. An examination of the PCI across both monkeys (0. 61 ± 0.12, t-test: *p* < 10^-6^) and humans (0.89 ± 0.1, t-test: *p* < 10^-6^) showed that, generally, subjects compensate significantly for the perturbations by appropriately adjusting their responses to reach the goal locations, with the humans’ PCI values being closer to ideal (**Fig. 3C**). Unlike monkeys, only three of the nine human subjects received end-of-trial feedback during this task (Methods). Nevertheless, compensation captured by PCI was comparable across both groups (with feedback: 0.86 ± 0.06, without feedback: 0.91 ± 0.12, t-test: *p* = 0.52).

In summary, the endpoints of subjects’ trajectories with perturbations are significantly different from those expected from an uncompensated behavior, demonstrating that both macaques and humans can integrate optic flow effectively. To better understand how subjects compensated for perturbations, we investigated the dynamic profile of the subjects’ responses as a function of the direction and magnitude of the perturbations. Representative example trials show different steering responses for forward versus backward perturbations. Subjects responded to backward perturbations by increasing their linear speed and extending travel duration (**Fig. 4A**, Trial 1&2). In contrast, subjects responded to forward perturbations by decreasing their speed and reducing their travel time (**Fig. 4A**, Trial 3&4). For the angular component, subjects rotated in the opposite direction to the perturbation’s angular velocity, even when the perturbation would have brought them closer to the target (**Fig. 4A**).

**Figure 4:**
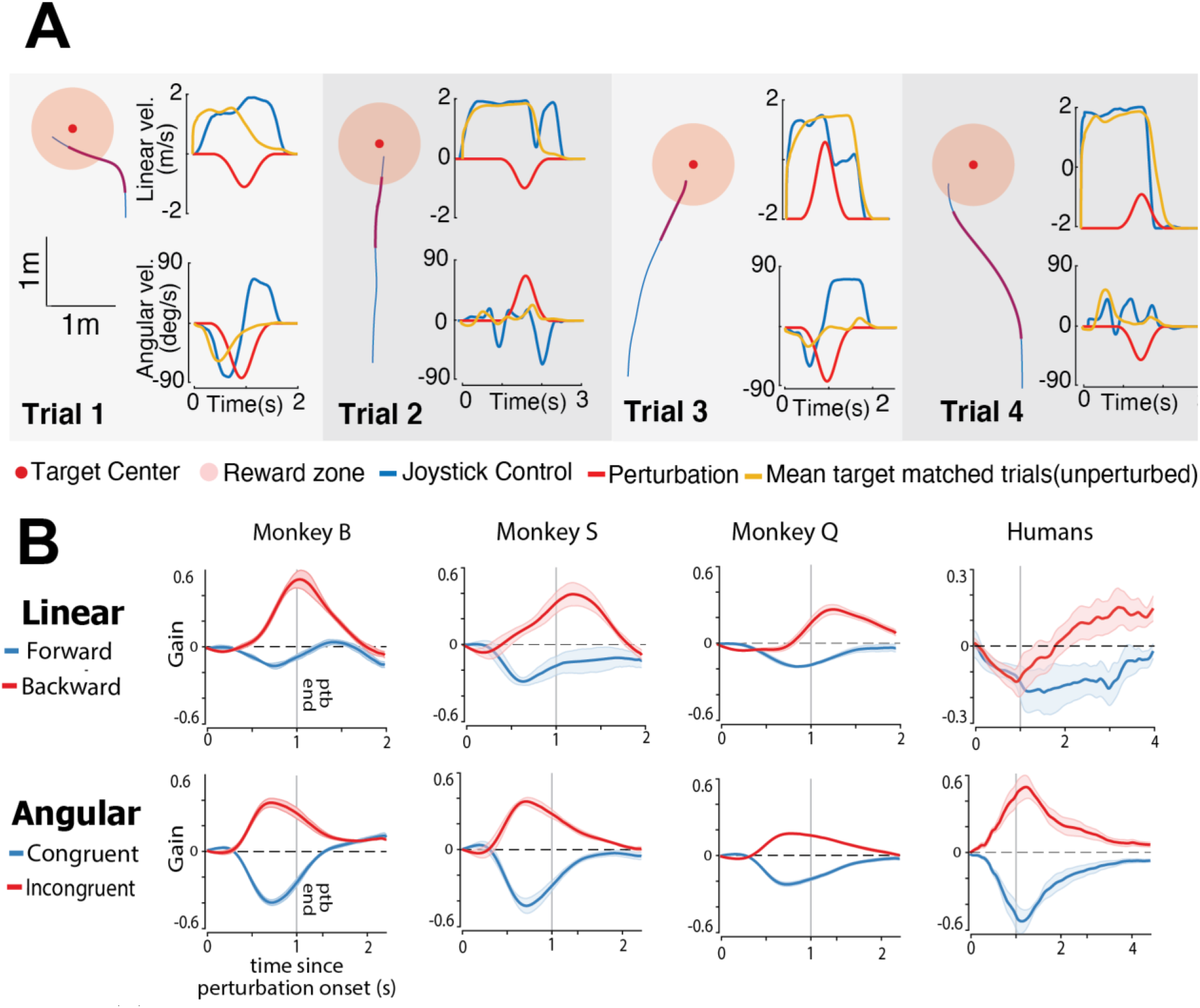
**(A) Left:** Aerial view of the trajectories for four trials with different perturbation profiles. Purple segments of the trajectories correspond to the perturbation period. **Right:** Time course for the linear (top) and angular (bottom) velocities. Red line: perturbation velocity profile; Blue line: joystick-controlled velocity of the current trial; Yellow line: mean trajectory for 50 target-matched trials without perturbations. **(B)** Average linear (*top*) and angular (*bottom*) components of individual monkey (*columns 1-3*) and human (*average across individuals*) responses to the perturbations, aligned to perturbation onset. Each component is grouped based on whether the perturbation drives the subject towards (blue: forward/congruent rotation) or away (red: backward/incongruent rotation) from the target. The response is normalized and expressed in units of the perturbation amplitude. Gray vertical line denotes the end of perturbation (Methods).

We grouped subjects’ responses based on whether the perturbation pushed subjects toward (forward/congruent) or away (backward/incongruent) from the target and computed average responses separately for the linear and angular perturbations (**Fig. 4B**; see **Methods**). Responses were estimated for each subject by computing the average deviation of self-motion during the perturbed trials from unperturbed target-matched trials, within a time window of 2s from the perturbation onset, normalized by the perturbation amplitude on that trial. The sign of the response denotes the direction of the response relative to the perturbation direction, and the amplitude indicates the strength of the compensation.

The amplitude of compensation was larger for perturbations that pushed monkeys backwards (away from the target) compared to those that pushed forwards (towards the target) (0.24 ± 0.11 (SD) vs. 0.44 ± 0.17, *t*-test: *p* = 0.002). In contrast, angular compensation was comparable for rotations towards (congruent) and away (incongruent) from the target (0.39 ± 0.14 vs. 0.34 ± 0.13, *p* = 0.31) (**Fig. 4B**). The reason for symmetric effects in the angular domain is that on most trials, monkeys were nearly done rotating towards the target by the time the perturbation arrived (example trials in **Fig. 4A**), and therefore did not really benefit from congruent perturbations. Qualitatively similar findings were seen for human subjects. For angular responses, humans rotated in the opposite direction of the perturbation’s angular velocity with response amplitudes that were similar for congruent and incongruent rotations (0.57 ± 0.15 vs. 0.56 ± 0.15, *p* = 0.84). For linear responses, humans were more conservative, as they slowed down at the time of perturbation onset (**Suppl. Fig. 1A**), and later producing the adequate response by increasing/decreasing their velocity or travel time depending on the perturbation direction. The response amplitude of human subjects was comparable for forward and backward perturbations (0.19 ± 0.08 vs. 0.16 ± 0.09, *p* = 0.64) **(Fig. 4B, Suppl. Fig. 1B)**. It is likely that the tendency of humans to slow down immediately after the perturbation onset allowed them to more effectively decouple the effects of self-motion and external perturbations on optic flow and compensate better than monkeys.

Nevertheless, despite these small differences between macaques and humans, these results indicate that subjects do use optic flow to dynamically adjust the speed and duration of steering according to the perturbation properties.

### Subjects adjust their velocity according to joystick control gain

In another version of the task, we manipulated the mapping between actions and their consequences by altering the gain of the joystick controller (**Fig. 5A**). In monkeys, the joystick control gain was altered to vary among 1x, 1.5x and 2x in separate blocks comprising 500 trials each. In humans, the gain factor varied randomly between 1x and 2x on different trials. To assess how much subjects adjust their responses to the different gain manipulations, we once again compared behavioral responses with hypothetical ‘uncompensated’ trajectories (**Fig. 5A**, dashed lines), computed by multiplying linear and angular responses during gain 1 x trials by the altered gain factor (1.5x or 2x). If subjects ignored the sensory feedback from optic flow cues, their steering would not be significantly different from the uncompensated responses.

**Figure 5.**
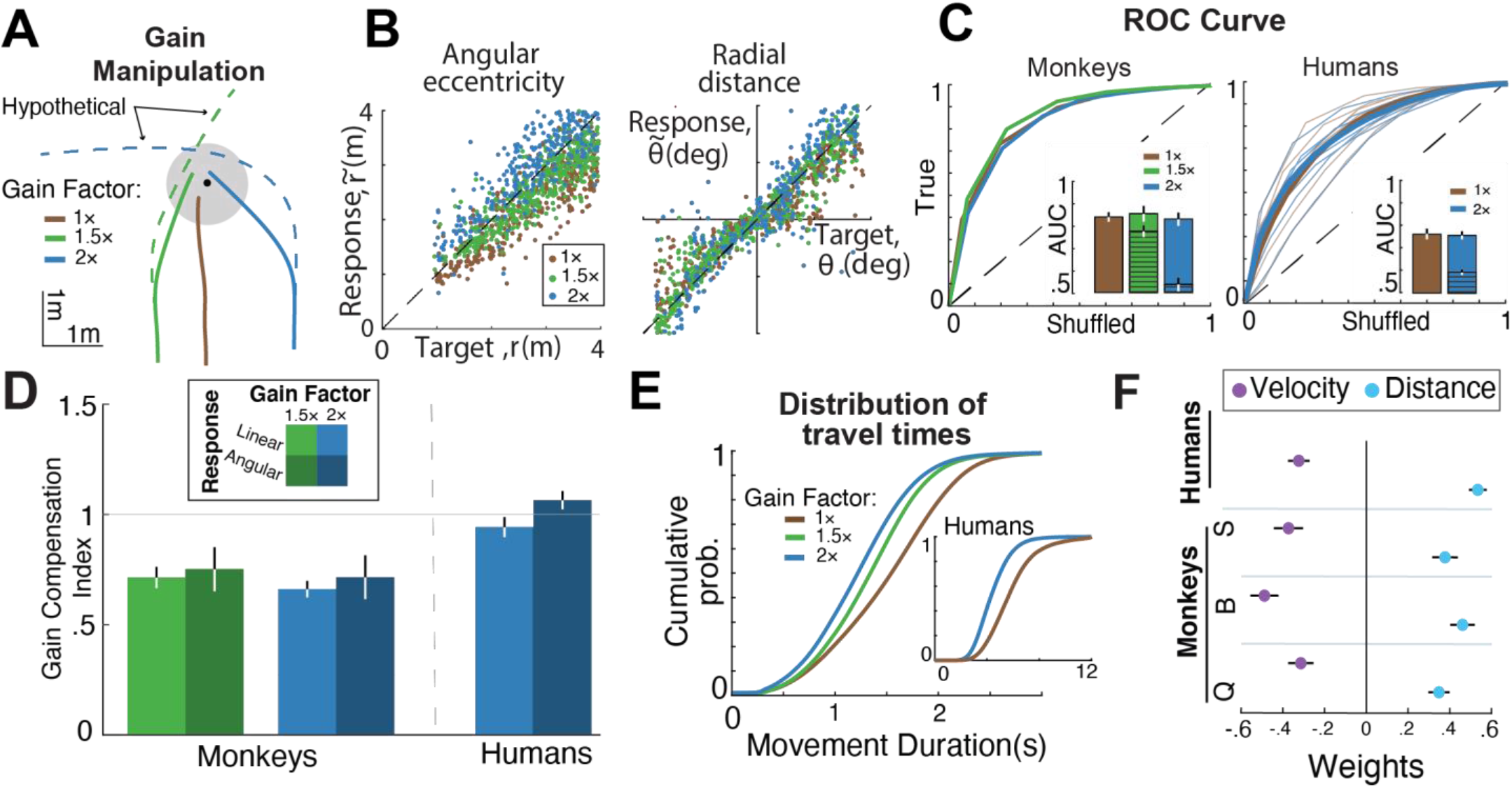
**(A)** Steering trajectories to an example target (black dot) with joystick gain factor x1 (solid brown curve), x1.5 (solid green curve) and x2 (solid blue curve). Dashed green and blue curves represent the hypothetical (uncompensated) trajectories with no compensation. **(B)** Radial (*left*) and angular (*right*) responses of an example monkey during trials from different gain conditions (brown: 1x, green: 1.5x, blue: 2x). **(C)** ROC curves of all monkeys (*left*) and humans (*right*) obtained by plotting proportion of correct trials against the corresponding chance-level proportion from shuffled data. Inset shows the area under the curve for real data (solid bars) and uncompensated trajectories (lined bars). **(D)** Bar plot of the mean gain compensation index for monkey and human subjects. Green and blue colors represent gain factor x1.5 and 2, respectively, and color saturation denotes response type (light color: linear, dark color: angular). Gray horizontal line denotes GCI value of 1. **(E)** Cumulative distribution of travel time for all monkeys (humans, inset) under different gain conditions. **(F)** Coefficients of regression model [log (*T*) = *w_r_*log (*r*) + *w_ν_*log (*ν*)] capturing the effects of gain and distance on travel time. Filled circles represent regression weights of distance (*cyan*) and gain (*purple*).

To test this, we computed multiplicative response biases by regressing the radial distance 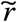 and angular eccentricity 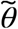 of the subjects’ final position against the initial target distance *r* and angle *θ* (**Fig. 5B**). For monkeys, both linear and angular biases increased with gain factor (mean ± standard deviation), Linear bias: 0.82 ± 0.03 (gain 1), 0.93 ± 0.08 (gain 1.5), 1.09 ± 0.08 (gain 2) (one way ANOVA: *p* < 10^-6^); Angular bias: 0.73 ± 0.05 (gain 1), 0.81 ± 0.12 (gain 1.5), 0.94 ± 0.24 (gain 2) (one-way ANOVA: *p* = 0.02)). Notably, in all cases, the animals’ biases were significantly smaller than the uncompensated response biases, suggesting that monkeys successfully compensated for changes in joystick gain (linear: paired *t*-test, *p* < 10^-6^(1.5x), *p* < 10^-6^ (2x); angular: *p* < 10^-3^ (1.5x), *p* < 10^-3^ (2x); **Suppl. Fig. 2**). Additionally, the area under the curve (**Fig. 5C**) was comparable for all joystick gains: 0.83 ± 0.03 (gain 1x), 0.86 ± 0.06 (gain 1.5x) and 0.82 ± 0.05 (gain 2x) (one way ANOVA: *p* = 0.12) and significantly different from the uncompensated case (gain 1.5x, *p* = 0.002; gain 2x, *p* < 10^-6^). Combined, these results suggest that monkeys adjust their steering according to the joystick control gain, further supporting the hypothesis that they take visual sensory input into account when choosing their actions.

Human subjects demonstrated even more ideal behavior. There was no difference in either the linear bias [mean ± standard deviation: 1.33 ± 0.29 (gain 1x) vs 1.27 ± 0.25 (gain 2x), *p* = 0.24] or the angular bias [1.6 ± 0.26 (gain 1x) vs 1.68 ± 0.28 (gain 2x), *p* = 0.23], yet bias differed from the uncompensated case responses (linear; *p* < 10^-3^, angular; *p* < 10^-3^). Similarly, there was no gain dependence in the measured AUC: 0.77 ± 0.06 (gain 1x) and 0.76 ± 0.06 (gain 2x) (*p* = 0.37), but both were higher than the uncompensated case (*p* < 10^-3^).

To further quantify subjects’ responses to the different control gains across humans and monkeys, we computed a gain compensation index (GCI). The GCI represents the extent of compensation relative to baseline (gain factor 1x, GCI of 0 means no compensation, GCI of 1 perfect compensation). Monkey GCIs averaged (± SE Linear/Angular) 0.71 ± 0.05/0.77 ± 0.09 (gain 1.5x) and 0.67 ± 0.03/0.72 ± 0.09 (gain 2x). Human GCIs were closer to ideal: 0.94 ± 0.04/1.06 ± 0.03 (gain 2x) (**Fig. 5D**).

Evidence for compensation was also seen in travel time. For humans, the mean travel time was lowest for highest gain: 4.48 ± 0.84s (gain 2x) vs. 5.89 ± 1.39 s (gain 1x) (paired t-test, *p* < 10^-3^). This was also true for monkeys (1.35 ± 0.17s (gain 1.5x), and 1.31 ± 0.19 (gain 2x) vs. 1.66 ± 0.23s (gain 1x) (one-way ANOVA, *p* < 10^-3^)). Thus, both humans and monkeys adapted to the different gain values by adjusting their travel duration appropriately (**Fig. 5E**).

Collectively, these results show that monkeys and humans adapt their responses to the changing joystick gain, supporting the hypothesis that subjects perform the task successfully by integrating optic flow. Even so, the sparse optic flow may not be the only information about the subjects’ spatial location in the virtual environment relative to the position of the target. Subjects could also partially incorporate predictions from an efference copy of their joystick movements.

To test the hypothesis that the subjects’ navigation strategy is based on optic flow integration, we contrasted it with pure time integration. To quantify the relative dependence on the two strategies, we took advantage of the lawful relationship between travel time, distance and velocity [velocity = distance/time]. Accordingly, we simultaneously regressed the travel time against initial target distance (*r*) and mean velocity (*ν*) in the log space across trials [log (*T*) = *w_r_*log (*r*) + *w_ν_*log (*ν*)] (see **Methods**). The travel time of an ideal path integrator would depend on changes in both distance and gain, with *w_r_* = 1 and *w_ν_* = −1. In contrast, the alternative strategy of pure time integration would predict weights *w_r_* = 1 and *w_ν_* = 0 (no dependence on velocity). Across all subjects, the regression weight on velocity, *w_ν_*, was significantly different from 0 in all monkey and human subjects ([95% confidence interval (CI) of regression weight], Monkey Q: [-0.34, −0.28], Monkey B: [-0.52, −0.46], Monkey S: [-0.41, −0.34], Humans: [-0.35, −0.30]) (**Fig. 5F**). The weight on target distance, wr, was positive and different from 0, Monkey Q: [0.32, 0.37], Monkey B: [0.43, 0.49], Monkey S: [0.34, 0.41], Humans: [0.51, 0.55]. (**Fig. 5F**). Notably, when this analysis was restricted to rewarded trials only, *w_ν_*, was closer to −1 (Monkey Q: [-0.99, −0.91], Monkey B: [-1.01, −0.92], Monkey S: [-1.02, −0.95], Humans: [-0.94, −0.88]), and *w_r_* was closer to 1 (Monkey Q: [0.84, 0.92], Monkey B: [0.83, 0.91], Monkey S: [0.84, 0.92], Humans: [0.94, −0.99]). This analysis of the responses to gain manipulation clearly supports the hypothesis that subjects perform the task by integrating optic flow.

### Optic Flow density affects task performance

A final manipulation involved changing the reliability of optic flow by varying the density of the ground plane elements between two possible values (“sparse” and “dense” for monkeys, “with” and “without” optic flow for humans; **Fig. 6A**). If subjects rely on optic flow integration to navigate, different values of ground plane density would impact the subjects’ responses. Indeed, the overall response variability was much larger for low density conditions across monkey subjects for both linear (standard deviation ± SE, high density: 0.56 ± 0.05 m, low density: 0.68 ± 0.05 m, t-test: *p* < 10^-3^) and angular (high density: 9 ± 0.8 deg, low density: 10 ± 0.8 deg, *p* < 10^-3^) responses (**Fig. 6B**). Likewise, in human subjects, the removal of optic flow increased the standard deviation of linear (with optic: 1.11 ± 0.08 m, w/o optic flow: 1.37 ± 0.07 m, t-test: *p* = 0.03) and angular (with optic flow: 14.15 ± 3.1deg, w/o optic low: 40.83 ± 3.3 deg, *p* = 0.01) responses. Altering the reliability of optic flow affected subjects’ accuracy by increasing the absolute error in monkeys (mean Euclidian error ± SD: high density: 0.65 ± 0.2m, low density: 0.8 ± 0.22 m, t-test *p* < 10^-6^) and humans (with optic flow: 1.68 ± 0.91m, w/o optic flow: 2.5 ± 0.73m, t-test: *p*= 0.003) (Note: all trials, not just those rewarded, were included). This difference was also reflected in ROC analysis (**Fig. 6C, D**; AUC ±SD, high density: 0.85 ± 0.06, low density: 0.79 ± 0.08, *p* < 10^-6^). Similarly, removal of optic flow cues in humans decreased AUC from 0.8 ± 0.09 (optic flow) to 0.67 ± 0.07 (no optic flow) (t-test: *p* = 0.012). Once again, these results collectively suggest that the subjects rely heavily on optic flow to navigate to the target.

**Figure 6:**
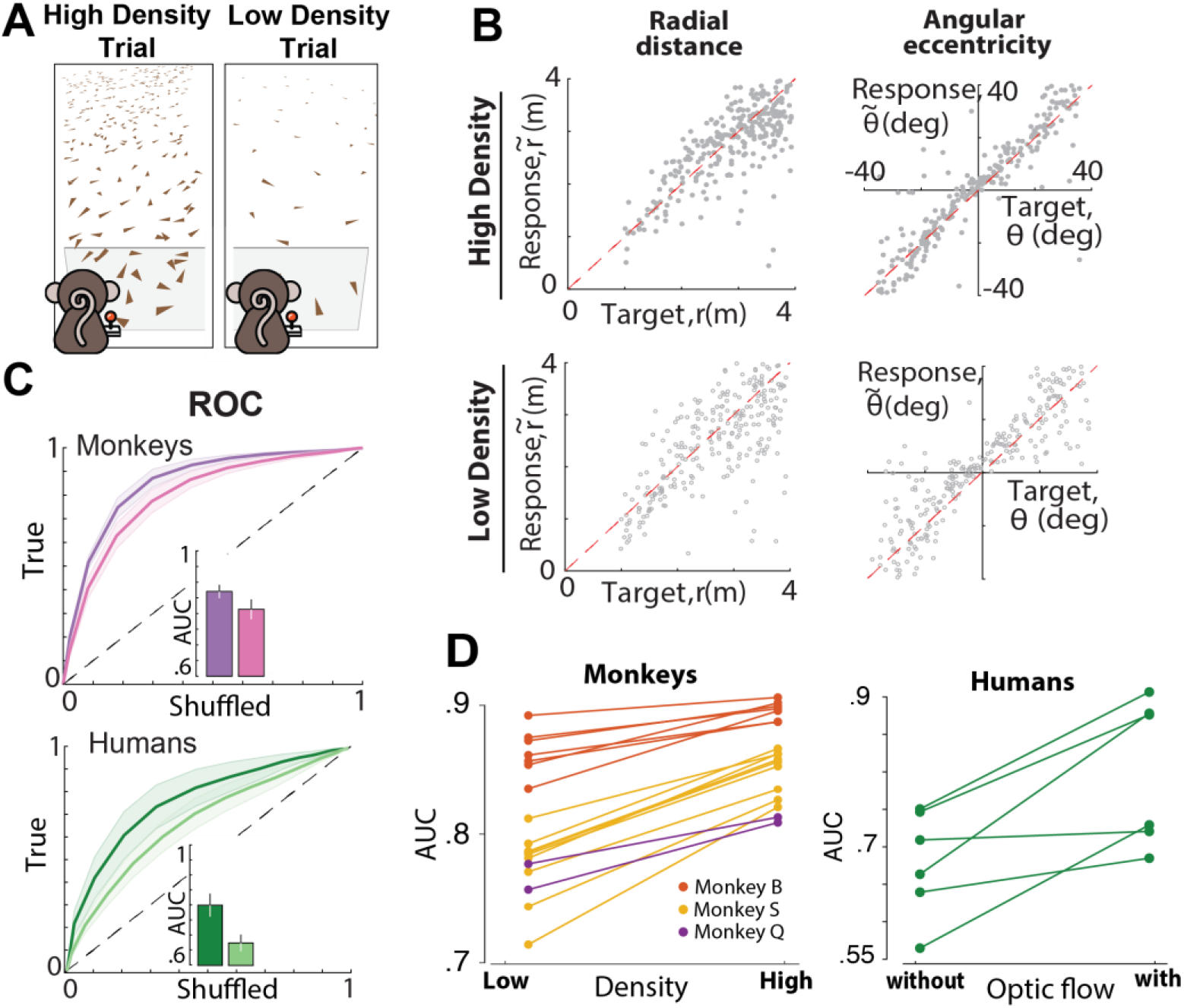
**(A)** Illustration of the two types of optic flow density conditions used throughout the experiment. The left and right sides of the graph denote trial conditions where ground elements occur at high and low densities respectively. **(B)** Radial and angular response of an example monkey with high (*top panels*) and low density (*bottom panels*) optic flow cues. **(C)** *Top:* ROC curves for all the monkeys for low (*pink*) and high (*purple*) density optic flow cues. *Bottom:* ROC curves for all the human subjects for trials with (*dark green*) and without (*optic flow*) optic flow cues. Shaded area represents SEM for all the human and monkey subjects respectively. Inset shows the area under the curve (AUC). **(D)** Pairwise comparison of the AUC for all the monkeys (left) and human subjects (right).

In a final variant of this task, we also manipulated the duration of the ground plane optic flow. Specifically, in humans (separate cohort, n=11) optic flow was presented for 500, 750, 1000, or 1500ms from trial onset (randomly intermixed across trials). We reasoned that if humans were not integrating optic flow across the duration of their trajectories (median duration ~2000ms), but instead were using the optic flow to initially adjust their internal models (e.g., their velocity estimate) and then navigated by dead reckoning, then their performance would not continuously improve with increasing optic flow durations. Performance, as measured by the area under the ROC, continuously improved with increasing optic flow durations (1-way ANOVA, F (3, 36) = 2.74, *p* < 0.05, Bonferroni-corrected post-hoc t-test, all *p* <0.05 see **Suppl. Fig. 3**), supporting the hypothesis that observers do indeed integrate optic flow over the entire duration of their trajectories.

In summary, we found that both macaques and humans were able to compensate for dynamic perturbations of internal state by using optic flow cues. In addition, subjects used optic flow to adjust their control and adapt to uncued changes in joystick gain. In both cases, the observed compensation was less than ideal, suggesting that subjects may also rely on an internal model of the control dynamics to some extent. Consistently with this, we found that removing optic flow cues decreased accuracy but did not completely blunt performance. Most critically, humans integrate optic flow throughout the duration of their trajectories.

## Discussion

Using three independent manipulations of the same basic navigation task, we showed that both monkey and human subjects rely on optic flow to navigate to the flashed target. Specifically, we introduced external perturbations to the subjects’ movement, varied the control gain of the joystick, and altered the density (and timing) of the optic flow to test whether the subjects integrate optic flow information to infer their self-location relative to the target. We found that subjects adjusted their steering velocity to compensate for external perturbations, adapted to joystick control gain changes, and showed degraded performance as the sensory uncertainty of the optic flow was decreased (monkeys) or eliminated (humans).

Human navigation research has long taken for granted that path integration by integration of optic flow is feasible, but never explicitly tested this hypothesis using naturalistic paradigms. Most previous studies used experimental paradigms that have artificial constraints, such as passive translation (Campos et al., 2012; Jürgens and Becker, 2006; Klatzky et al., 1998; Petzschner and Glasauer, 2011; Tramper and Medendorp, 2015), discretized decision and actions (Horst et al., 2015; Chrastil et al., 2016; Koppen et al., 2019), or restricted movements to a one dimensional track (Campbell et al., 2018; Frenz and Lappe, 2005; Frenz et al., 2007). In contrast, real-world navigation is an active process with rich temporal dynamics that typically takes place in two dimensions. By incorporating all three features into the task, our findings validate the utility of optic flow for navigation in the absence of landmarks.

The approach used here also has implications for studying sensory evidence accumulation in general. Traditionally, evidence accumulation has been studied using paradigms in which subjects passively view a noisy stimulus with predefined dynamics, integrate evidence for a period of time, and then report their decision at the end of the trial (Gold and Shadlen, 2007; Kiani et al., 2013). In such tasks, the latent dynamics are not under the subject’s control. The task used in this study offers a way to study evidence accumulation in the more naturalistic setting where latent dynamics (and thus the sensory inputs) are constantly influenced by the subject’s actions. This is important because decisions in the real world are often governed both by sensory inputs as well as expectations from one’s own actions. Although rodent decision-making research has started moving in this direction by training animals to accumulate evidence while navigating in a T-maze (Pinto et al., 2018), the range of possible actions in such tasks is still limited. In contrast, subjects here could use both linear and angular components of optic flow to navigate towards a continuum of possible target locations. The moderate complexity of this task can more accurately capture real-world dynamics and allow for future exploration of sensory evidence accumulation and decision-making with fewer restrictions than traditional binary tasks, without sacrificing the ability to manipulate task variables(Noel et al., 2021).

Our findings support an optic flow-based navigation strategy that is conserved across humans and monkeys, thus paving way for the study of neural mechanisms of path integration in monkeys. There is already a well-documented hierarchy of regions in the macaque brain that are involved in processing optic flow (Britten, 2008), including a representation of both linear and angular velocity in the posterior parietal cortex (Avila et al., 2019). Analyzing the relationship of neural responses in those areas to the animal’s estimates during binary decision tasks have helped better the understanding of the feedforward mechanisms underlying heading perception (Lakshminarasimhan et al., 2018b; Pitkow et al., 2015). Analyzing neural recordings during richer tasks such as the one used here calls for more sophisticated tools (Balzani et al., 2020; Kwon et al., 2020; Wu et al., 2020), but will likely shed light on the recurrent mechanisms that underlie more dynamic computations of perceptual decision making in closed-loop dynamic tasks.

It must be noted that results from all three manipulations were consistent with a strategy that combines sensory feedback control based on optic flow cues, and predictive control based on an internal model of the dynamics. Such a combination has been extensively reported in the context of motor control tasks (Wolpert and Ghahramani, 2000; Wolpert et al., 1995). Thus, similar principles may underlie control of limb movements and visuomotor control using external affordances, such as driving a car. Given the rich experimental evidence for the role of cerebellum in constructing internal models (Ito, 2008), cerebellar targets to the posterior parietal cortex (Bostan et al., 2013) may prove important in the fine between internal model predictions and sensory feedback signals such as optic flow.

## Methods

### EQUIPMENT AND TASK

Three rhesus macaques (all male, 7-8 yrs. old) participated in the experiments. All surgeries and experimental procedures were approved by the Institutional Animal Care and Use Committee and were in accordance with National Institutes of Health guidelines.

Additionally, four distinct groups of human subjects participated in the three variants of the experiment: nine human subjects (6 males, 3 females, age: 20-30) in the perturbation variant, seven (4 males, 3 females, age:18-30) in the gain variant, six (4 males, 2 females, age: 20-30) in the density variant, and eleven (6 males, 5 females, age: 21-30) in the time-varying density variant. All human subjects were unaware of the purpose of the study and signed an approved consent form prior to their participation in the experiment.

### Experimental setup

At the beginning of each experimental session, monkeys were head-fixed and secured in a primate chair placed on top of a platform (Kollmorgen, Radford, VA, USA). A 3-chip DLP projector (Christie Digital Mirage 2000, Cypress, CA, USA) was mounted on top of the platform and rear-projected images onto a 60 × 60 cm tangent screen ~30cm in front of the monkey. The projector was capable of rendering stereoscopic images generated by an OpenGL accelerator board (Nvidia Quadro FX 3000G). Spike2 software (Power 1401 MkII data acquisition system from Cambridge Electronic Design Ltd.) was used to record joystick and all event and behavioral markers for offline analysis at a sampling rate of 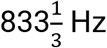.

All stimuli were generated and rendered using C++ Open Graphics Library (OpenGL) by continuously repositioning the camera based on joystick inputs to update the visual scene at 60 Hz. The virtual camera was positioned at a height of 10cm above the ground plane. Spike2 software (Power 1401 MkII data acquisition system from Cambridge Electronic Design Ltd.) was used to record and store the target location (*r, θ*), subject’s position 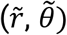.

For humans, all other aspects of the setup were similar to the one used for monkeys, but with subjects seated 67.5cm in front of a 149 × 127 cm^2^ (width × height) rectangular screen, and with the virtual camera placed 100cm above the ground plane. The time-varying optic flow density manipulation was presented in virtual reality (HTC VIVE, New York, NY) and built in Unity (San Francisco, CA).

### Behavioral task

Subjects used an analog joystick (M20U9T-N82, CTI electronics) with two degrees of freedom and a square displacement boundary to control their linear and angular speed in a virtual environment. This virtual world was comprised of a ground plane whose textural elements had a limited lifetime (~250ms) to avoid serving as landmarks. The ground plane was circular with a large radius of 70m (near and far clipping planes at 5cm and 4000cm respectively), and the subject was positioned at its center at the beginning of each trial. Each texture element was an isosceles triangle (base × height: 8.5 × 18.5 cm^2^) which was randomly repositioned and reoriented anywhere in the arena at the end of its lifetime, making it impossible to use as a landmark. The maximum linear and angular speeds were initially fixed to *ν*_max_ = 2m^-1^ and *ω*_max_ = 90°/s, respectively, and then varied by a factor of 1.5 and/or 2.0. The density of the ground plane was either held fixed at *ρ* = 2.5 elements/m^2^ or varied randomly between two values (*ρ* = 2.5 elements/m^2^ and *ρ* = 0.1 elements/m^2^) in a subset of recording sessions (see below). The stimulus was rendered as a red-green anaglyph and projected onto the screen in front of the subject’s eyes. Except for when wearing a virtual reality headset, subjects wore goggles fitted with Kodak Wratten filters (red #29 and green #61) to view the stimulus. The binocular crosstalk for the green and red channels was 1.7% and 2.3% respectively. Target positions were uniformly distributed within the subjects’ field of view with radial distances and angles that varied from 1 to 4 m and −35 to 35 degrees respectively for monkey experiments. In the human experiments, the radial distance and the angle of the targets varied from 1 to 6 m and −40 to 40 degrees, respectively. In the time-varying optic flow experiment in humans the targets varied from −30 to 30 degrees in eccentricity but were always presented at 3 m in radial distance.

Monkeys received binary feedback at the end of each trial. They either received a drop of juice if, after stopping, they were within 0.6m away from the center of the target; otherwise, no juice was provided. The fixed reward boundary of 0.6m was determined using a staircase procedure prior to the experiment to ensure that monkeys received reward in approximately two-thirds of the trials. Human subjects did not receive feedback during the experiment, with the exception of 3 subjects, for which feedback consisted of a bull’s eye pattern consisting of six concentric circles (with the radius of the outermost circle being continuously scaled up or down by 5%, according to the one-up, two-down staircase procedure), displayed with an arrowhead indicating the target location on the virtual ground. The arrowhead presented was displayed either in red or green to denote whether the participant’s response had occurred within the outermost rewarded circle (See Lakshminarasimhan et al., 2020 for details).

### Movement Dynamics

Let *s_t_*, *o_t_* and *a_t_* denote the participant’s state (velocity), observation (optic flow), and action (joystick position) respectively. The equations governing movement dynamics in this experiment are as follows:

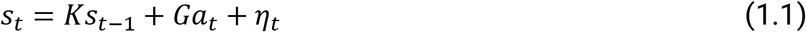

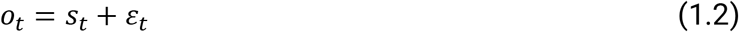

where *K* denotes resistance to change in state, *G* denotes the gain factor of the joystick controller, *η_t_* denotes process noise, and *ε_t_* denotes observation noise. In all our experiments, we set *K* = 0 such that there was no inertia. Participants can navigate by integrating their velocity to update position *x* as *x*_*t*+1_ = *x_t_* + *s_t_*Δ*t* where Δ*t* denotes the temporal resolution of updates. Note that the experiment involved two degrees of freedom (linear and angular), so the above equation applies to both.

### Behavioral manipulations

In the perturbation variant of the experiment, normal trials were interleaved with trials that incorporated transient optic flow perturbations to dislocate subjects from their intended trajectory. Mathematically, such perturbations can be understood as setting the process noise (*η_t_* in Equation 1.1) to a nonzero value. For <u>monkeys,</u> the perturbations had a fixed duration of 1s, a velocity with a Gaussian profile of σ = .2 and an amplitude drawn from a uniform distribution from −2 to 2m/s, and from −120 to 120 deg/s for the linear and angular velocities, respectively. For monkeys, the perturbations had a fixed duration of 1s, a Gaussian velocity profile with σ = .2, and an amplitude drawn from a uniform distribution between −2 and 2m/s and −120 and 120°/s for the linear and angular velocities, respectively. For <u>humans,</u> the perturbations had also a fixed duration of 1s, whereas their velocity profile was an isosceles triangle with height that varied with a uniform distribution from −2 to 2m/s, and −120 to 120°/s for the linear and angular velocities, respectively. For both humans and monkeys, the perturbation onset time was randomly varied from 0 to 1s after movement onset.

In the gain manipulation variant, we switched the gain factor of the joystick controller (*G* in Equation 1.1) among 1, 1.5, or 2 for monkeys, and between 1 and 2 for humans. For monkeys, we manipulated joystick control in separate blocks of 500 trials, and the ordering of the blocks was randomized between days. For humans, the gain factor varied randomly between trials. Within each trial, both linear and angular velocities were scaled by the same gain factor.

In the density manipulation variant for monkeys, the density of ground plane elements varied between two values, “high” (2.5 elements/m^2^) and “low” (0.1 elements/m^2^). For humans, normal trials were interleaved with trials where the elements constituting the ground plane were completely removed. Density manipulation effectively changes the magnitude of observation noise (*ε_t_* in Equation 1.2) and is a way of controlling sensory reliability.

### Data Analyses

Customized MATLAB code was written to analyze data and to fit models. Depending on the quantity estimated, we report statistical dispersions either using 95% confidence interval, standard deviation, or standard error of the mean. The specific dispersion measure is identified in the portion of the text accompanying the estimates. For error bars in figures, we provide this information in the caption of the corresponding figure.

Across all animals and humans, we regressed (without an intercept term) each subject’s response positions 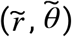 against target positions (*r, θ*) separately for the radial (*r* vs 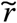) and angular (*θ* vs 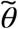) coordinates, and the radial and angular multiplicative biases were quantified as the slope of the respective regressions. The 95% confidence intervals were computed by bootstrapping.

### Simulation of Uncompensated Case (Perturbations)

To simulate the uncompensated case responses for each trial with a perturbation, we picked an unperturbed trial with the most similar target position to the perturbed trial. Then, we added the angular and linear perturbation velocities (preserving their timing) to the steering angular and linear velocities of the monkey to simulate an ‘uncompensated’ stopping position if there was no compensation. Specifically, for each time step *t* within the window of the perturbation duration, we added the instantaneous linear and angular components of the perturbation *α_t_* and *β_t_* from the perturbed trial to the linear and angular steering velocity *ν_t_* and *ω_t_* of the chosen target-matched unperturbed trial, respectively. As a result, the total linear 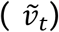 and angular 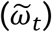 instantaneous velocities of the trial during the simulated perturbation were:

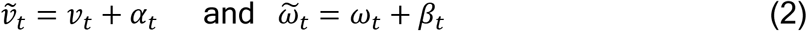

The velocity time series of the whole trial were then integrated to produce a ‘uncompensated’ response with no compensation.

### ROC Analysis

To quantify and compare subject performance across conditions (unperturbed, perturbations, uncompensated case), we performed ROC analysis as follows. For each subject, we first calculated the proportion of correct trials as a function of a (hypothetical) reward boundary. In keeping with the range of target distances used, we gradually increased the reward boundary until reaching a value that would include all responses. Whereas an infinitesimally small boundary will result in all trials being classified as incorrect, a large enough reward boundary will yield near-perfect accuracy. To define a chance-level performance, we repeated the above procedure, this time by shuffling the target locations across trials, thereby destroying the relationship between target and response locations. Finally, we obtained the ROC curve by plotting the proportion of correct trials in the original dataset (true positives) against the shuffled dataset (false positives) for each value of hypothetical reward boundary. The area under this ROC curve was used to obtain an accuracy measure for all the subjects.

### Perturbation Compensation Index (Perturbations)

To quantify the subjects’ compensatory responses to the perturbations, we computed the Perturbation Compensation Index (PCI) based on the results of the ROC analysis:

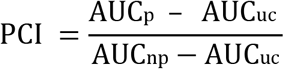

where AUC_p_ represents the area under the curve (AUC) for trials with perturbations, AUC_uc_ is the AUC for the group of the uncompensated case responses, and AUC_np_ is the AUC for the unperturbed trials. A value of 0 indicates an accuracy equal to the uncompensated case response and thus represents a complete lack of compensation, whereas a value of 1 indicates an accuracy equivalent to the unperturbed trial and thus represents perfect compensation.

### Response to perturbations

For each subject, we estimated a perturbation-specific response by computing the trial-averaged deviation in the subject’s self-motion velocity (relative to target-matched, unperturbed trials) at various time lags between 0-2s from perturbation onset, in steps of 6ms. The rationale behind computing the *deviation* from target-matched unperturbed trials rather than *raw* velocities is that we essentially subtract the component of the subject’s response that is influenced by target location, yielding only the perturbation-specific component. This deviation is normalized by the perturbation amplitude before trial averaging, such that the sign denotes the direction of the response relative to the direction of perturbation, and the amplitude denotes the strength of compensation.

### Gain Compensation Index (Gain Manipulation)

To quantify the subjects’ responses for different joystick gain conditions, we computed the gain compensation index (GCI), which measures the extent to which the subjects compensated for the changes in gain factor (*g* = [1.5, 2]) with respect to the baseline gain (*g_0_* = 1):

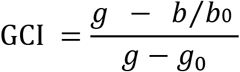

where *g* and *g*_0_ denote the modified gain factor and the baseline gain factor respectively. *b* and *b*_0_ correspond to the multiplicative bias (radial or angular) of the block of trials with the modified gain factor and baseline, respectively. The ratio *b/b*_0_ captures the change in multiplicative bias for the block of trials where the joystick gain was modified. A ratio equal to 1 denotes a perfect compensation, whereas a value of 0 denotes a complete lack of compensation.

### Multiple Linear Regression Model (Gain Manipulation)

In order to test whether subjects perform spatial (as opposed to temporal) integration, we expressed the basic kinematic equation of velocity: *ν* = *x/t* in the form log(*t*) = log(*x*) - log (*ν*) which allowed for the implementation of a multiple linear regression. Following earlier work (Kwon and Knill, 2013), we assume that noise variance is constant in logarithmic scale. To measure the influence of distance and velocity we used the model:

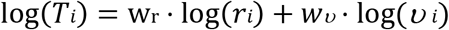

where T, r and *ν* are travel duration, target distance and mean velocity of trial *i*, respectively.

## Data and software availability

MATLAB code implementing all quantitative analyses in this study is available online (https://https://github.com/panosalef/fireflyTask). Datasets generated by this study are available online (https://gin.g-node.org/panosalef/sensory_evidence_accumulation_optic_flow).

## Acknowledgements

We thank Jing Lin and Jian Chen for their technical support, and Baptiste Caziot and Akis Stavropoulos for their useful insights. This work was supported by the NIH (1U19-NS118246 - BRAIN Initiative, 1R01 DC014678).

## Supplemental Materials

**Suppl. Table 1.** shows Pearson correlation coefficients ± s.d. between monkeys’ and humans’ responses and target locations (separately for radial and angular components). Correlation coefficients were computed separately across the set of unperturbed (left), perturbed (middle) and hypothetical uncompensated (right) trials. *p*-values between columns indicate the significance of *t*-test for the difference between the correlation correlations shown in the surrounding columns.

|  |  | Unperturbed | p | Perturbed | p | Uncompensated |
| --- | --- | --- | --- | --- | --- | --- |
| Monkeys | Angular | 0.91 ± 0.03 | 4·10<sup>-10</sup> | 0.85 ± 0.04 | 2.9·10<sup>-13</sup> | 0.76 ± 0.05 |
|  | Radial | 0.73 ± 0.1 | 1.4·10<sup>-15</sup> | 0.59 ± 0.08 | 4·10<sup>-3</sup> | 0.33 ± 0.08 |
| Humans | Angular | 0.94 ± 0.03 | 1.3·10<sup>-12</sup> | 0.89± 0.04 | 10<sup>-45</sup> | 0.62 ± 0.05 |
|  | Radial | 0.38 ± 0.05 | 2.6·10<sup>-5</sup> | 0.33± 0.1 | 0.9 | 0.33 ± 0.08 |

**Suppl. Figure 1:**
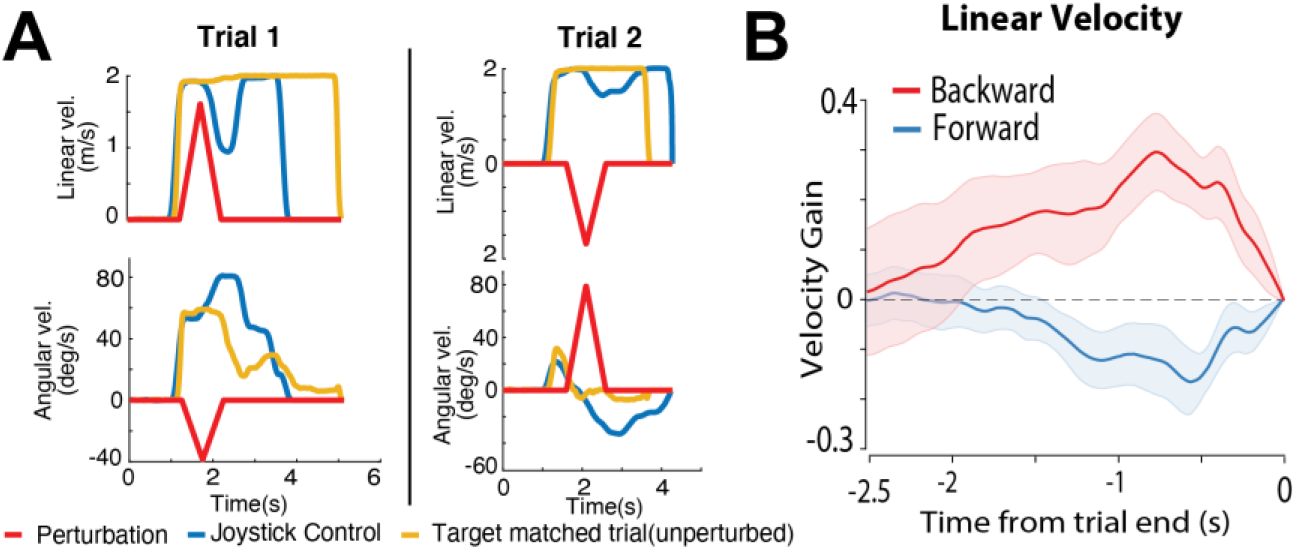
**(A)** Time course for the linear (top) and angular (bottom) velocities of two example trials with perturbations with a human subject. Red: perturbation velocity profile; blue: joystick-controlled velocity of the current trial; yellow: trajectory of a target-matched trial without perturbations. **(B)** Because subjects typically try to reach the target quickly by traveling at highest speeds, the rational method to compensate for backward/forward perturbations is by traveling for longer/shorter duration (than they would’ve in the absence of perturbation). Because the average target distance, and thus travel duration, was longer in human experiments, the timing of this compensation could be delayed with respect to the perturbation onset and thus not fully captured by the average responses within a small window aligned to perturbation onset (Figure 4B). For this reason, we computed a response of the linear velocity component for forward (*blue*) and backward (*red*) perturbation aligned to the trial end. Responses are computed relative to target-matched trials (see *methods*) and demonstrate that subjects decrease their travel times in order to compensate for the forward perturbations as shown by the negative deflection of the response (*blue line*), whereas they increase their travel time for backward perturbations as shown from the positive deflection (*red line*).

**Suppl. Figure 2.**
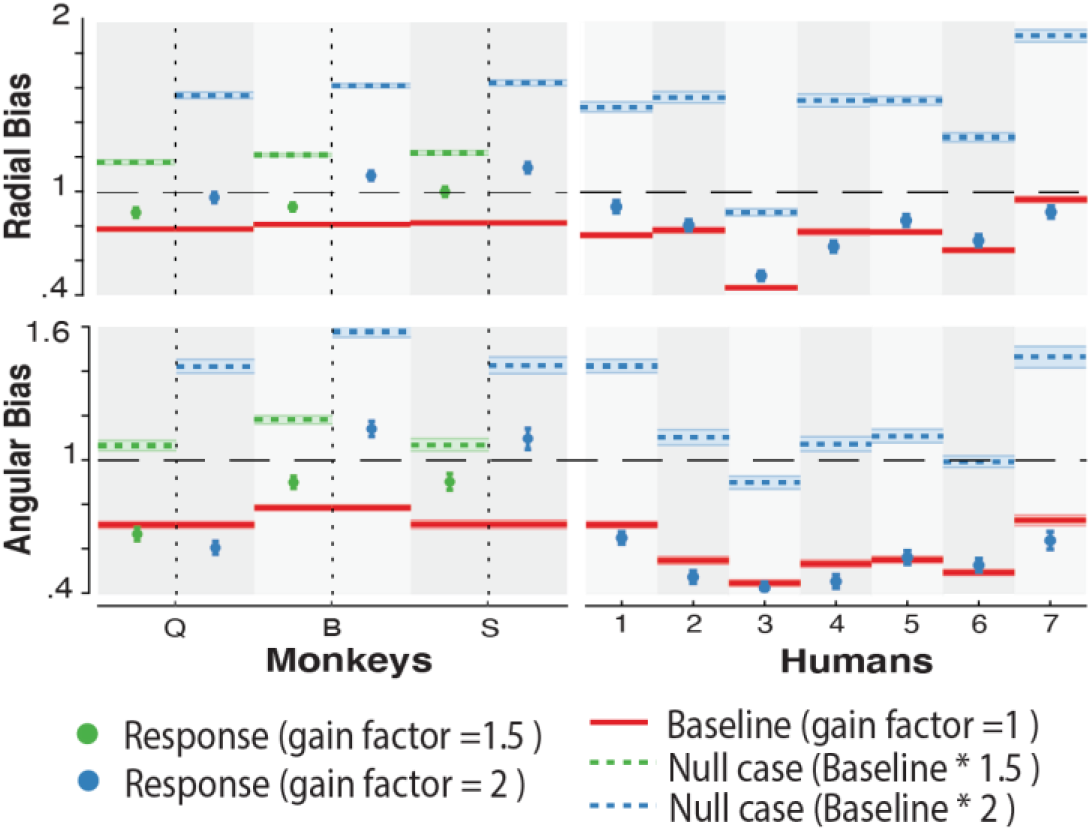
Radial (top) and Angular (bottom) Bias for all monkeys and humans. Red lines represent bias for gain factor = 1(baseline bias), dashed lines represent the uncompensated case, defined as the baseline bias multiplied by the gain factor of the corresponding block of trials, and dots represent the real bias of the subjects.

**Suppl. Figure 3:**
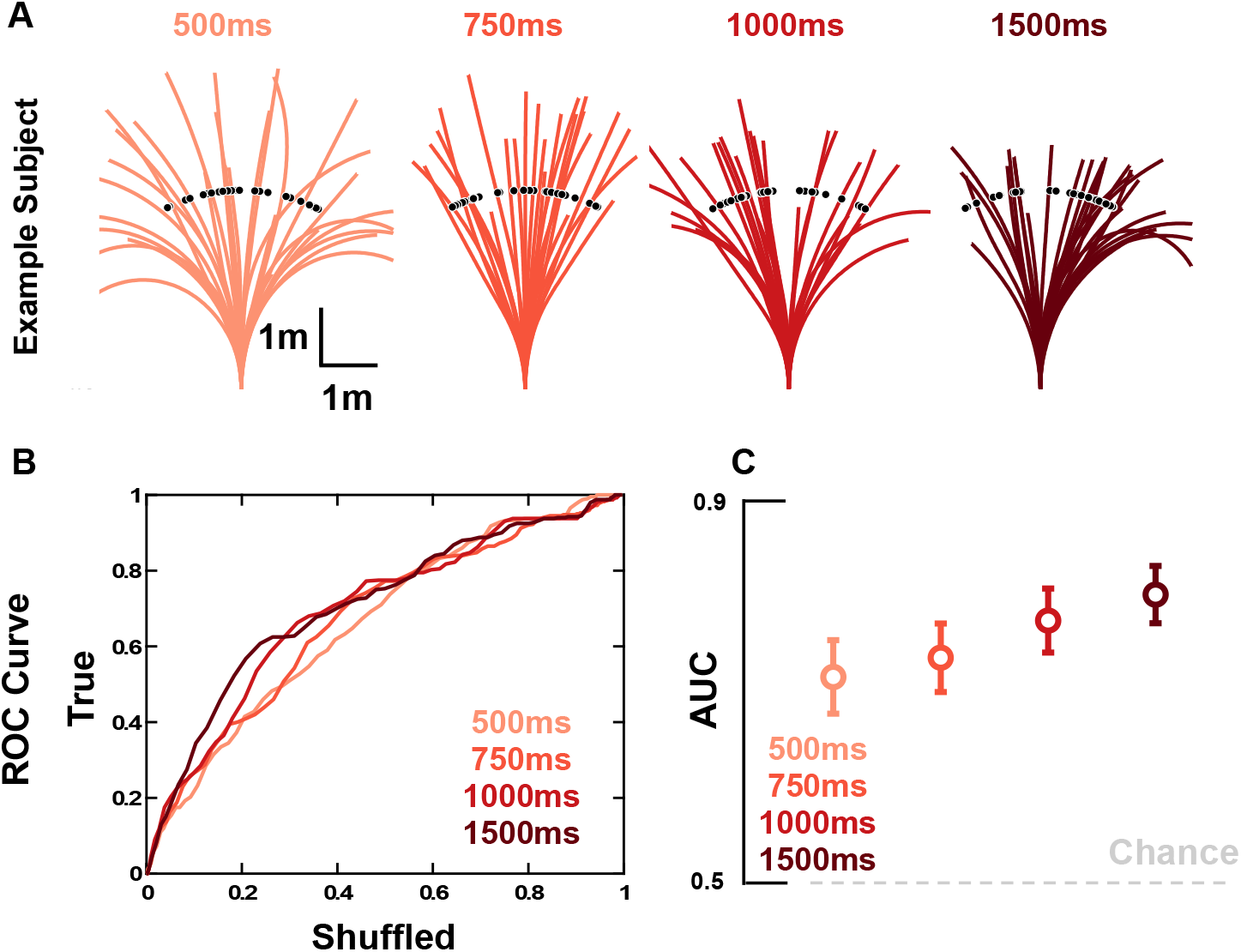
**(A)** Firefly locations (black dots) and trajectories for a subset of trials in an example subject. As the presentation of optic flow information increased in duration (red gradient, from left to right: 500, 750, 1000, and 1500ms) the subject’s overshooting of targets became less drastic. This is in line with a known prior for slow speeds in humans during this task (Lakshminarasimhan et al., 2018; Noel et al., 2020), given that at lower optic flow presentation durations the trajectory ought to be further driven by the prior. **(B)** ROC curves for all subjects (thin line) and their average (thick line) as a function of optic flow duration. **(C)** Mean area under the curve (AUC) of the ROC. Error bars are +/- 1 S.E.M.

## References

Åström, K.J. (1965). Optimal control of Markov processes with incomplete state information. J. Math. Anal. Appl.

Avila, E., Lakshminarasimhan, K.J., DeAngelis, G.C., and Angelaki, D.E. (2019). Visual and Vestibular Selectivity for Self-Motion in Macaque Posterior Parietal Area 7a. Cereb. Cortex 29, 3932–3947.

Balzani, E., Lakshminarasimhan, K., Angelaki, D.E., and Savin, C. (2020). Efficient estimation of neural tuning during naturalistic behavior. 34.

Bostan, A.C., Dum, R.P., and Strick, P.L. (2013). Cerebellar networks with the cerebral cortex and basal ganglia. Trends Cogn. Sci.

Britten, K.H. (2008). Mechanisms of Self-Motion Perception. Annu. Rev. Neurosci.

de Bruyn, B., and Orban, G.A. (1988). Human velocity and direction discrimination measured with random dot patterns. Vision Res.

Butler, J.S., Smith, S.T., Campos, J.L., and Bülthoff, H.H. (2010). Bayesian integration of visual and vestibular signals for heading. J. Vis.

Campbell, M.G., Ocko, S.A., Mallory, C.S., Low, I.I.C., Ganguli, S., and Giocomo, L.M. (2018). Principles governing the integration of landmark and self-motion cues in entorhinal cortical codes for navigation. Nat. Neurosci.

Campos, J.L., Butler, J.S., and Bülthoff, H.H. (2012). Multisensory integration in the estimation of walked distances. Exp. Brain Res.

Collett, M., and Collett, T.S. (2017). Path Integration: Combining Optic Flow with Compass Orientation. Curr. Biol.

Collett, T.S., and Collett, M. (2000). Path integration in insects. Curr. Opin. Neurobiol. 10, 757–762.

Drugowitsch, J., Deangelis, G.C., Angelaki, D.E., and Pouget, A. (2015). Tuning the speed-accuracy trade-off to maximize reward rate in multisensory decision-making. Elife.

Ellmore, T.M., and McNaughton, B.L. (2004). Human path integration by optic flow. Spat. Cogn. Comput.

Etienne, A.S., and Jeffery, K.J. (2004). Path integration in mammals. Hippocampus.

Frenz, H., and Lappe, M. (2005). Absolute travel distance from optic flow. Vision Res.

Frenz, H., Bührmann, T., Lappe, M., and Kolesnik, M. (2007). Estimation of Travel Distance from Visual Motion in Virtual Environments. ACM Trans. Appl. Percept.

Glass, L., and Pérez, R. (1973). Perception of random dot interference patterns. Nature.

Gold, J.I., and Shadlen, M.N. (2000). Representation of a perceptual decision in developing oculomotor commands. Nature.

Gold, J.I., and Shadlen, M.N. (2007). The Neural Basis of Decision Making. Annu. Rev. Neurosci.

Golledge, R.G. (1999). Human navigation by path integration. Wayfinding Behav. Cogn. Mapp. Other Satial Process. 125–151.

Gu, Y., Angelaki, D.E., and DeAngelis, G.C. (2008). Neural correlates of multisensory cue integration in macaque MSTd. Nat. Neurosci.

Heinze, S., Narendra, A., and Cheung, A. (2018). Principles of Insect Path Integration. Curr. Biol.

Hou, H., Zheng, Q., Zhao, Y., Pouget, A., and Gu, Y. (2018). Neural Correlates of Optimal Multisensory Decision Making. BioRxiv.

Ito, M. (2008). Control of mental activities by internal models in the cerebellum. Nat. Rev. Neurosci.

Jürgens, R., and Becker, W. (2006). Perception of angular displacement without landmarks: Evidence for Bayesian fusion of vestibular, optokinetic, podokinesthetic, and cognitive information. Exp. Brain Res.

Kautzky, M., and Thurley, K. (2016). Estimation of self-motion duration and distance in rodents. R. Soc. Open Sci.

Kearns, M.J., Warren, W.H., Duchon, A.P., and Tarr, M.J. (2002). Path integration from optic flow and body senses in a homing task. Perception.

Kiani, R., Churchland, A.K., and Shadlen, M.N. (2013). Integration of direction cues is invariant to the temporal gap between them. J. Neurosci. 33, 16483–16489.

Kim, J.N., and Shadlen, M.N. (1999). Neural correlates of a decision in the dorsolateral prefrontal cortex of the macaque. Nat. Neurosci.

Klatzky, R.L., Loomis, J.M., Beall, A.C., Chance, S., and Golledge, R.G. (1998). Spatial updating of selfposition and orientation during real. Psychol. Sci.

Kwon, O.S., and Knill, D.C. (2013). The brain uses adaptive internal models of scene statistics for sensorimotor estimation and planning. Proc. Natl. Acad. Sci. U. S. A.

Kwon, M., Schrater, P., Daptardar, S., and Pitkow, X. (2020). Inverse rational control with partially observable continuous nonlinear dynamics. ArXiv.

de Lafuente, V., Jazayeri, M., and Shadlen, M.N. (2015). Representation of accumulating evidence for a decision in two parietal areas. J. Neurosci.

Lakshminarasimhan, K.J., Petsalis, M., Park, H., DeAngelis, G.C., Pitkow, X., and Angelaki, D.E. (2018a). A Dynamic Bayesian Observer Model Reveals Origins of Bias in Visual Path Integration. Neuron.

Lakshminarasimhan, K.J., Pouget, A., DeAngelis, G.C., Angelaki, D.E., and Pitkow, X. (2018b). Inferring decoding strategies for multiple correlated neural populations. PLoS Comput. Biol. 14.

Lakshminarasimhan, K.J., Avila, E., Neyhart, E., DeAngelis, G.C., Pitkow, X., and Angelaki, D.E. (2020). Tracking the Mind’s Eye: Primate Gaze Behavior during Virtual Visuomotor Navigation Reflects Belief Dynamics. Neuron.

Lappe, M., Jenkin, M., and Harris, L.R. (2007). Travel distance estimation from visual motion by leaky path integration. Exp. Brain Res.

Liu, T., and Pleskac, T.J. (2011). Neural correlates of evidence accumulation in a perceptual decision task. J. Neurophysiol.

Noel, J.P., Lakshminarasimhan, K.J., Park, H., and Angelaki, D.E. (2020). Increased variability but intact integration during visual navigation in Autism Spectrum Disorder. Proc. Natl. Acad. Sci. U. S. A.

Noel, J.P., Caziot, B., Bruni, S., Fitzgerald, N.E., Avila, E., and Angelaki, D.E. (2021). Supporting generalization in non-human primate behavior by tapping into structural knowledge: Examples from sensorimotor mappings, inference, and decision-making. Prog. Neurobiol. 201, 101996.

Petzschner, F.H., and Glasauer, S. (2011). Iterative Bayesian estimation as an explanation for range and regression effects: A study on human path integration. J. Neurosci.

Pinto, L., Koay, S.A., Engelhard, B., Yoon, A.M., Deverett, B., Thiberge, S.Y., Witten, I.B., Tank, D.W., and Brody, C.D. (2018). An accumulation-of-evidence task using visual pulses for mice navigating in virtual reality. Front. Behav. Neurosci.

Pitkow, X., Liu, S., Angelaki, D.E., DeAngelis, G.C., and Pouget, A. (2015). How Can Single Sensory Neurons Predict Behavior? Neuron 87, 411–423.

Snowden, R.J., and Braddick, O.J. (1990). Differences in the processing of short-range apparent motion at small and large displacements. Vision Res.

Stone, T.J. (2017). Mechanisms of place recognition and path integration based on the insect visual system.

Sutton, R.S., and Barto, A.G. (1998). Reinforcement Learning: An Introduction. IEEE Trans. Neural Networks.

Thurley, K., and Ayaz, A. (2017). Virtual reality systems for rodents. Curr. Zool.

Tramper, J.J., and Medendorp, W.P. (2015). Parallel updating and weighting of multiple spatial maps for visual stability during whole body motion. J. Neurophysiol.

Watanabe, O., and Kikuchi, M. (2006). Hierarchical integration of individual motions in locally paired-dot stimuli. Vision Res.

Wiener, M., Michaelis, K., and Thompson, J.C. (2016). Functional correlates of likelihood and prior representations in a virtual distance task. Hum. Brain Mapp.

Wolpert, D.M., and Ghahramani, Z. (2000). Computational principles of movement neuroscience. Nat. Neurosci.

Wolpert, D.M., Ghahramani, Z., and Jordan, M.I. (1995). An internal model for sensorimotor integration. Science (80-.).

Wu, Z., Kwon, M., Daptardar, S., Schrater, P., and Pitkow, X. (2020). Rational thoughts in neural codes. Proc. Natl. Acad. Sci. U. S. A.

